# Multiscale entropy analysis of combined EEG-fNIRS measurement in preterm neonates

**DOI:** 10.1101/2023.07.12.548724

**Authors:** Lorenzo Semeia, Mina Nourhashemi, Mahdi Mahmoudzadeh, Fabrice Wallois, Katrin Sippel, Hubert Preissl

## Abstract

In nature, biological systems such as the human brain are characterized by complex and non-linear dynamics. One way of quantifying signal complexity is Multiscale Entropy (MSE), which is suitable for structures with long-range correlation at different time scales. In developmental neuroscience, MSE can be taken as an index of brain maturation, and can differentiate between healthy and pathological development. In our current work, we explored the developmental trends of MSE on the basis of 30 simultaneous EEG – fNIRS recordings in premature infants between 27 and 34 weeks of gestational age (wGA). To explore potential factors impacting MSE, we determined the relation between MSE and the EEG Power Spectrum Density (PSD) and Spontaneous Activity Transients (SATs). As a result, via wGA, the MSE calculated on the EEG increases, thus reflecting the maturational processes in the brain networks, whereas in the fNIRS, MSE decreases, which might indicate a maturation of the brain blood supply. Moreover, we propose that the EEG power in the beta band (13-30 Hz) might be the main contributor to MSE in the EEG. Finally, we highlight the importance of SATs in determining MSE as calculated from the fNIRS recordings.

**Highlights:** Biological systems show complex and non-linear dynamics. With Multiscale Entropy (MSE) we studied simultaneous EEG-fNIRS in premature infants. MSE in the EEG increases over gestational age, MSE in the fNIRS decreases. EEG power spectrum density and spontaneous activity transients contribute to MSE.

## 1. Introduction

According to the World Health Organization, in the year 2018, around 15 million children were born prematurely, i.e., before the 37^th^ week of gestation. In comparison to infants born at term, these babies also have a higher risk of impaired neurological development, and monitoring their brain activity is therefore essential (Benders et al., 2015; El-Dib et al., 2022; Saigal & Doyle, 2008).

In addition, the development of the neurovascular system in this population is not fully understood. One of the most common techniques for monitoring brain function in preterm infants is resting-state electroencephalography (EEG) which is well-suited for neonatal intensive care units. The features of premature EEG vary considerably from week to week between the 26^th^ and 40^th^ week of gestational age (wGA) (O’Toole, Boylan, Vanhatalo, & Stevenson, 2016; Wallois et al., 2021). This reflects the progressive increase in cortical volume (Garcia et al., 2018), connectivity (Stoecklein et al., 2020), neural migration, molecular specification, myelination, and receptors maturation (e.g. Kostovic & Rakic, 1990; Kostović, 2020). In addition, the brain activity in premature infants results from different neural dynamics. Successive spontaneous endogenous oscillations, for example, are specific to functions (e.g., theta temporal activities in coalescence with slow waves (TTA-SW), delta brushes and frontal sharp transients). Furthermore, the progressive establishment of the neuronal network involves the progressive decrease in discontinuity, visible in the time-domain, and the progressive increase in the dominant frequency power (Wallois et al., 2021). The neural activity discontinuity consists of quiescent periods separated by bursts of activities such as the spontaneous oscillations nestling in ultraslow oscillations known as spontaneous activity transients (SATs). SATs are believed to be particularly relevant for neural wiring and networks definition in the developing brain (Ackman, Burbridge, & Crair M.C., 2012; Arichi et al., 2017; Feller, 1999; Omidvarnia, Fransson, Metsäranta, & Vanhatalo, 2014). Beginning at around 28 wGA, with the establishment of thalamic afferents to the cortical plate, the neural network also increases its dependency on experience (Kostovic & Rakic, 1990; Kostović, 2020). Finally, during this period in which the premature neonate approaches its full term, the differentiation between sleep state emerges (Bourel-Ponchel, Hasaerts, Challamel, & Lamblin, 2021). A premature birth affects this sleep states organization in a way that might carry prognostic information (Tokariev et al., 2019; Yrjölä, Stjerna, Palva, Vanhatalo, & Tokariev, 2022). Given the connection between neural and vascular systems that actively constitute the neurovascular coupling (NVC), SATs provide us with an opportunity to study the spontaneous hemodynamic response to neural activation even without external stimulation (Nourhashemi, Mahmoudzadeh, Goudjil, Kongolo, & Wallois, 2020; Roche-Labarbe, Wallois, Ponchel, Kongolo, & Grebe, 2007).

In adults, the NVC is well defined, and consists of an increase in blood flow in the surroundings of the neural activation (Iadecola, 2017; Jöbsis, 1977). The NVC begins early during gestation and can be detected in preterm neonates at 28 weeks of gestational age (Roche-Labarbe et al., 2008). Although otherwise similar to adults, the NVC in neonates and infants has age-dependent features (Arichi et al., 2012). In a recent study, the characteristics of NVC in preterm infants between 27 and 35 weeks of gestational age (GA) at rest were investigated using EEG and functional near-infrared spectroscopy (fNIRS) (Nourhashemi et al., 2020). In particular, the group focused on the coupling between the spontaneous bursts of neural activity and the hemodynamic patterns. Their results highlight how the NVC – in the same preterm infant and during the same recording – could take on three different forms in response to similar neural activity. In addition, these age-related coupling features in the NVC differ from those in adults.

Complex and non-linear dynamics characterize features of many biological systems found in nature, including the human brain. Complexity measurement of brain signals delivered promising results in the study of human brain during spontaneous activity, of cognition, of disorders such as Parkinson, epilepsy, anesthesia, autism, Alzheimer, and of the human brain during development (Anokhin, Lutzenberger, Nikolaev, & Birbaumer, 2000; Hadoush, Alafeef, & Abdulhay, 2019; Lutzenberger, Birbaumer, Flor, Rockstroh, & Elbert, 1992; Lutzenberger, Flor, & Birbaumer, 1997; Müller, Lutzenberger, Preißl, Pulvermüller, & Birbaumer, 2003; Müller, Lutzenberger, Pulvermüller, Mohr, & Birbaumer, 2001; Rodriguez-Bermudez, G., & Garcia-Laencina, P. J., 2015; Sun et al., 2020). Physiological data can be well characterized by multiscale entropy (MSE (Costa, Goldberger, & Peng, 2005) which is suited for structures with long-range correlations on different time scales. In developmental neuroscience, MSE can be used as an index for brain maturation and showed the highest overall usability, comparability to other works in literature, and computational efficiency in comparison to other metrics (e.g., Moser et al., 2019). In particular, MSE of EEG signals shows a linear increase with age from birth to adulthood (McIntosh, Kovacevic, & Itier, 2008); (Lippé, Kovacevic, & McIntosh, 2009; van Noordt & Willoughby, 2021). Wel et al. (Wel et al., 2017) recently measured MSE in resting state EEG of preterm infants born before 42 weeks of GA, and reported a positive correlation between MSE and GA between 27 and 42 weeks of GA. In addition, they used MSE to differentiate between neonatal sleep stages. Quiet sleep in particular is associated with a lower complexity index than active sleep.

Signal complexity calculated from fNIRS signals in preterm infants is a rather recent but promising approach. For example, da Sortica Costa et al. (Da Sortica Costa et al., 2017) described how a reduced NIRS signal MSE was associated with brain injury and mortality. Moreover, transfer entropy between EEG and fNIRS signals was lower in preterm neonates with brain abnormalities than in healthy neonates (Hendrikx et al., 2020).

Overall, multiscale entropy constitute a suitable tool for the study of brain activity, especially considering its ability to capture multiple time scales, its relative easy interpretability and resistance to physiological noise, and its relation to brain functional connectivity (Liu, Song, Liang, Knöpfel, & Zhou, 2019; Wang et al., 2018).

Regarding brain maturation, however, we are not aware of studies that investigate brain complexity in preterm infants using a multimodal approach that combines simultaneous electroencephalography (EEG) and functional near-infrared spectroscopy (fNIRS) recordings. In this work, we explore MSE in simultaneous EEG – fNIRS signals of preterm infants to evaluate its age-related changes and relate the complexity between neural and hemodynamic signals. The analysis of MSE is performed on the dataset from Nourhashemi et al. (2020). Moreover, we test whether power spectral density (PSD) and/or SATs are related to MSE. We hypothesize that these are the two major contributors to MSE. In fact, the discontinuous and non-stationarity of EEG signal in infants is characterized by SATs which can be changed with age by occurrence, duration and amplitude, and which can impact MSE calculation.

Moreover, since entropy is sensitive to spectral power, the MSE results with regard to the correlation between PSD and MSE in adults should be interpreted with caution (Kosciessa, Kloosterman, & Garrett, 2020). After **1)** determining the development of MSE, PSD and SAT over time in simultaneous EEG – fNIRS recordings in preterm infants between 27 and 35 weeks of GA, we **2)** explore the relationship between MSE, PSD, SATs and GA by performing a stepwise regression. This serves to detect which factors might be related to or determine MSE at a premature age. Finally, **3)** the relation between these variables is further explored by mediation analysis.

## 2. Methods

### 2.1. Participants, recordings and datasets

The participants’ particulars, recordings and datasets are described in Nourhashemi et al. (2020). In short, the study consists of 32 preterm neonates (12 females, with a mean gestational age (GA) of 29.3 weeks at birth, and ranging between 27 and 34 weeks GA), who were recorded in a supine position (mean GA at test of 31.3 weeks). The ethical committee of the Amiens University Hospital approved this study (CPP Nord-Ouest II-France IDRCB-2008-A00704-51). All the parents gave their informed consent in accordance with the Declaration of Helsinki, and agreed to their data being used for further research. The neurophysiological recordings were performed using the simultaneous, multimodal approach as described in Nourhashemi et al. (Nourhashemi et al., 2020). This includes EEG and Continuous Wave (CW) fNIRS. The multimodal recording probe was constructed so as to be placed on the infant’s forehead to record the spontaneous physiological data (for further details on the recording system, see Nourhaschemi et al. (2020)). For the current analysis, we included one frontal EEG channel out of the eight available, corresponding to Fp1 in the international 10-20 system. The choice of channel was simplified by the fact that this was the only channel where EEG and fNIRS sensors were placed on the same spatial position. The datasets length varied between 12 and 55 minutes, with a mean of 32 and standard deviation of 10 minutes. The EEG sampling frequency was of 1024Hz. The fNIRS sampling frequency was 5Hz. Before preprocessing, two datasets were excluded since no simultaneous EEG – fNIRS recording was possible due to technical problems, resulting in N = 30 datasets included for further analyses. The whole analysis was performed with Matlab R2019b for Windows.

### 2.2. Preprocessing

EEG datasets were initially linearly detrended. Data were then band-pass filtered between 1 and 30 Hz. After filtering, data were cleaned from high amplitude artefacts using the artefact blocking (AB) algorithm (Mourad, Reilly, de Bruin, Hasey, & MacCrimmon, 2007). In the AB algorithm, a threshold was calculated for each participant, corresponding to the median value of the data time-series plus three standard deviations of the signal. Finally, data were downsampled to 256Hz.

With the CW-fNIRS, we acquired the concentration changes in oxygenated hemoglobin (HbO), de-oxygenated hemoglobin (HbR), and calculated the cerebral Tissue Oxygenation Index (TOI). In addition and offline, we calculated the difference between HbO and HbR, which is a proxy for oxygen consumption (HbD). These data were also initially detrended with a linear fit. The datasets were next band-pass filtered between 0.03 and 0.5 Hz following the example by Nourhashemi et al. (2020). The AB algorithm was also applied (Mourad et al., 2007). The threshold for the AB algorithm was calculated as for the EEG data.

### 2.3. Parameters estimation

Before proceeding with the estimation of our parameters of interest, we divided the EEG datasets into non-overlapping one-minute windows, resulting in N = 15360 data points per-window. We divided the datasets in non-overlapping windows of N = 2000 data points in the fNIRS signal. This number was selected to ensure that there were enough data-windows for each dataset, and that these sufficed to perform MSE analysis up to scale 20 (Angelini et al., 2007; Da Sortica Costa et al., 2017). The MSE and PSD were calculated for each of the windows defined in the EEG, HbO, HbR, HbD and TOI signal. Outliers were identified using the generalized “ESD many-outliers” procedure (Rosner, 1983) and removed using the algorithm by Taliaferro (2021). The values of these parameters were averaged across windows within subject. Despite the fact that dividing a dataset into separate windows decreases the data-points for the parameters’ calculation, this procedure is more proficient in reducing noise and artefacts that are common in neurophysiological recordings in newborns. SAT detection (see paragraph below) was applied to the EEG dataset as a whole, but not to the single data-windows. A list of the parameters calculated is depicted in *Table 1*.

**Table 1.**
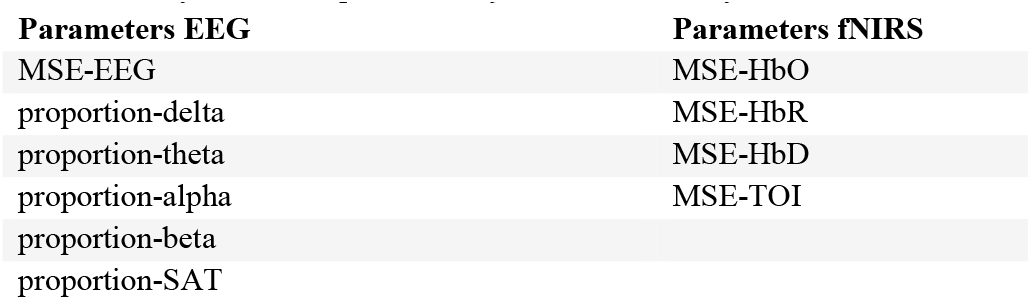
List of calculated parameters for both EEG and fNIRS.

| **Parameters EEG** | **Parameters fNIRS** |
| --- | --- |
| MSE-EEG | MSE-HbO |
| proportion-delta | MSE-HbR |
| proportion-theta | MSE-HbD |
| proportion-alpha | MSE-TOI |
| proportion-beta |  |
| proportion-SAT |  |

#### 2.3.1. Multiscale Entropy

MSE was calculated by the ‘msentropy’ function implemented in the WFDB toolbox for the analysis of physiological data, which is available in Matlab (Goldberger et al., 2000; Silva & Moody, 2014). For the calculation of scales 1-20, we used all the default parameters. The WFDB toolbox uses for MSE computation the codes from Costa, Goldberger, & Peng (2002) and Costa, Goldberger, & Peng (2005). The frequencies in the signals covered by our complexity scales were calculated by dividing the sampling frequency by the scale length. The EEG was processed with a sampling frequency of 256 Hz and scales 2-20, resulting in a frequency interval between 12.8 and 128 Hz. For the sampling frequency of 5 Hz in the fNIRS device, the frequency range was between 0.25 and 2.5 Hz. Due to the use of a band pass filter (0.03 and 0.5 Hz), this resulted in a complexity analysis in the interval 0.25 – 0.5 Hz. Further analysis was based on the following parameters: MSE-EEG, MSE-HbO, MSE-HbR, MSE-HbD, MSE-TOI.

#### 2.3.2. Spectral analysis

In the EEG data, we calculated PSD using the Fast Fourier Transform algorithm (‘fft’) implemented in Matlab. We determined the mean power in four frequency ranges, namely delta (1-4 Hz), theta (4-8 Hz), alpha (8-13 Hz) and beta (13-30 Hz). Finally, we calculated the proportions of PSD in each frequency band in relation to the overall power.

We therefore had four parameters for further analysis: proportion-delta, proportion-theta, proportion-alpha, proportion-beta.

#### 2.3.3. SAT detection

SATs in the EEG recordings were detected using a Non-Linear Energy Operator (NLEO) (O’Toole, Temko, & Stevenson, 2014). We then defined proportion-SAT as the proportion of data-points detected as SAT over the total duration of the dataset. An example of SAT detected in the EEG is depicted in *Figure 1a*.

**Figure 1.**
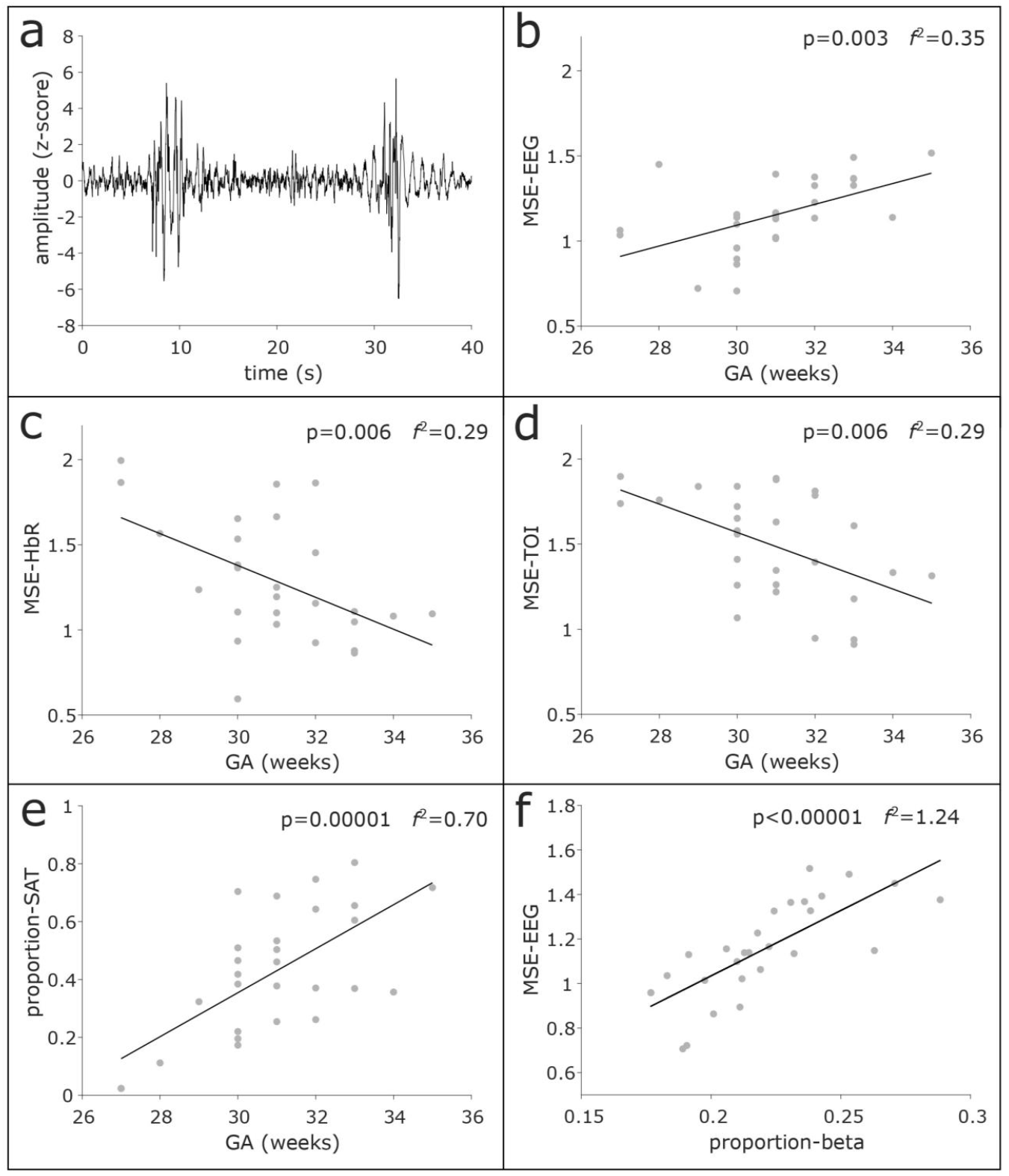
a) an example of EEG recording in a preterm neonate displaying SAT; b) increase of MSE-EEG during GA; c) decrease of MSE-HbR during GA; d) decrease of MSE-TOI during gestation; e) positive linear relation between the proportion of data-points detected as SAT and GA; f) Correlation between MSE-EEG and proportion-beta. The higher the proportion-beta, the higher the MSE-EEG.

### 2.4. Statistics

We further checked and removed outliers applying a generalized extreme Studentized deviate test, with significance level at 0.05 and 15 maximum points tested (Taliaferro, 2021). Finally, the data were tested for normal distribution with the Kolmogorov-Smirnov test before proceeding with the statistical testing.

#### 2.4.1. Step 1. Linear models

To detect the development of different complexity metrics over gestation, linear models were built for each of the metrics, and values of p < 0.05 were considered statistically significant. The model is:

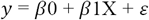

where *y* is one of the variables (MSE-EEG, MSE-HbO, MSE-HbR, MSE-HbD, MSE-TOI, proportion-delta, proportion-theta, proportion-alpha, proportion-beta and proportion-SAT), *β*0 is the intercept, *β*1 the regression coefficient, X the gestational age, and *ε*the error in the estimate. Effect size, defined by Cohen’s *f*^*2*^, is also taken into consideration. We applied Holm correction for multiple comparisons considering N=10 possible p-values.

#### 2.4.2. Step 2. Stepwise regressions

We investigated the relationship between MSE, PSD, SATs and GA by performing a stepwise regression analysis in which one of the MSE variables (MSE-EEG, MSE-HbO, MSE-HbR, MSE-HbD and MSE-TOI) is taken as the dependent variable. The other variables, such as the proportion-delta, proportion-theta, proportion-alpha, proportion-beta, proportion-SAT, GA and the MSE parameters which are not taken as dependent variable are included as possible predictors. We used the “stepwiselm” function in Matlab with default parameters to calculate the stepwise regression, where terms are added or removed iteratively. The default starting model is ‘constant’, which begins with only the constant term. The function evaluates available terms and adds the best one to the model if its associated F-test has a p-value of 0.05 or less. If no terms can be added, it examines the existing model and removes the least significant term if its F-test has a p-value of 0.10 or greater. This iterative process continues until no further additions or removals are possible. Notably, the constant term is always retained. The function may introduce two-way interactions among the predictors.

#### 2.4.3. Step 3. Mediation analysis

We also performed a mediation analysis to further investigate the relationship between MSE, PSD, SATs, and GA. This involved the use of the MediationToolbox (Tor Wager, 2022) implemented in Matlab. GA was considered as predictor, one of the five MSE parameters (MSE-EEG, MSE-HbO, MSE-HbR, MSE-HbD, MSE-TOI) as outcome variable, and one of the proportion-delta, proportion-theta, proportion-alpha, proportion-beta, proportion-SAT as mediator factor. With five mediation models for each of the five MSE parameters, a total of 25 mediation models were calculated. We applied Holm correction for multiple comparisons considering N=20 possible p-values. This number results from multiplying the number of p-values of each model (4) for the number of possible mediator factors for each of the MSE parameters (5).

## 3. Results

### 3.1. Step 1. Linear models

The linear models between MSE-EEG, MSE-HbO, MSE-HbR, MSE-HbD, MSE-TOI and GA revealed a developmental trend of these parameters during gestation. In particular, MSE-EEG increases during GA (p = 0.003, *f*^*2*^ = 0.35, *Fig. 1b*). The fNIRS-derived parameters, such as MSE-HbR (p = 0.006, *f*^*2*^ = 0.29) and MSE-TOI (p = 0.006, *f*^*2*^ = 0.29) decreased during GA (*Fig 1c and Fig 1d*, respectively). MSE-HbO (p = 0.01, *f*^*2*^ = 0.25) and MSE-HbD (p = 0.01, *f*^*2*^ = 0.23) also tended to decrease, but corresponding p-values did not survive correction for multiple comparisons (*Supplementary figure 1*).

The linear models between proportion-delta, proportion-theta, proportion-alpha, proportion-beta and GA also revealed some developmental trends of these parameters during gestation, even if did not survived correction for multiple comparisons (*supplementary figure 2*). In particular, proportion-alpha (p = 0.008, *f*^*2*^ = 0.27) tended to increase during GA, while there where no changes during gestation for proportion-beta (p = 0.06), proportion-delta (p = 0.1), and proportion-theta (p=0.09).

We also observed a positive linear relation between proportion-SAT and GA (p = 0.0001, *f*^*2*^ = 0.70, *Figure 1e*).

### 3.2. Step 2. Stepwise regressions

The linear models resulting from the stepwise regression analysis are reported in *Table 2*. Specifically, GA and proportion-beta are the main predictors for MSE-EEG (*R*^*2*^=0.65, p<0.001), MSE-HbR and MSE-HbD predict MSE-HbO (*R*^*2*^=0.0.86, p<0.001), proportion-SAT and MSE-HbO predict HbR (*R*^*2*^=0.85, p<0.001), MSE-HbO predict MSE-HbD (*R*^*2*^=0.70, p<0.001), and proportion-alpha, MSE-HbO, MSE-HbD and MSE-EEG predict MSE-TOI (*R*^*2*^=0.76, p<0.001).

**Table 2.**
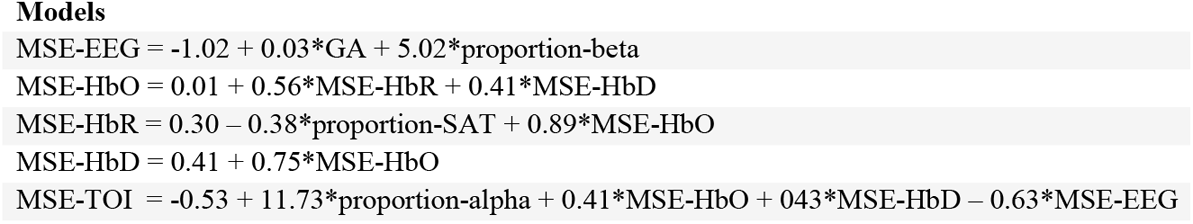
Stepwise regression models for MSE calculated for EEG and fNIRS recordings.

### 3.3. Step 3. Mediation analyses

Mediation analyses were computed to determine whether proportion-SAT, proportion-delta, proportion-theta, proportion-alpha, or proportion-beta are mediators in the association between GA (weeks) and MSE-EEG, MSE-HbO, MSE-HbR, MSE-HbD, and MSE-TOI. When MSE-EEG was taken as an outcome variable (*Supplementary figure 3*), the analysis revealed no mediation between GA (weeks) and MSE-EEG by any of the possible mediators, nor did we find mediation between GA (weeks) and MSE-HbO by our possible mediators when MSE-HbO was taken as the outcome variable (*supplementary figure 4*). Note that the proportion-beta is strongly related to MSE-EEG (p<0.00001). The relationship between these two parameters was further explored, and results are depicted in *Figure 1f*. Similar results were found for MSE-HbR (*Supplementary figure 5*), MSE-HbD (*Supplementary figure 6*), and MSE-TOI (*Supplementary figure 7*).

## 4. Discussion

In this study, we investigated signal complexity, quantified with MSE, of simultaneous EEG and fNIRS signals in a group of preterm infants between 27 and 34 weeks of GA. We also related MSE to other physiological parameters such as the power spectral density of the neural signal and the occurrence of SAT. We identified a linear relation between MSE calculated for both the EEG and the fNIRS signal and GA. Moreover, PSD and occurrence of SAT also displayed changes during GA which are in line with previous reports in the literature. From our stepwise regression and mediation analyses, we determined that the proportion of beta power is the major contributor to EEG signal complexity.

Moreover, the proportion of time detected as SAT is related to the MSE in the deoxygenated hemoglobin signal, highlighting how bursts of activity might influence MSE calculation in fNIRS data.

### 4.1. Development of signal MSE in preterm infants

We reported an increase of MSE-EEG signal of preterm infants during GA (*Figure 1b*), which is in line with previous reports (Niemarkt et al., 2011; Wel et al., 2017). Specifically, an increase over age of EEG signal complexity has been related to age-dependent maturation of endogenous brain networks (Janjarasjitt, Scher, & Loparo, 2008a; van Noordt & Willoughby, 2021). This is due to both structural and functional maturational changes. On the structural level, myelination, synaptic reorganization, cell differentiation, and relocation of thalamic afferents from the subplate to the cortical plate are examples of such dynamics (e.g. Kostovic & Rakic, 1990; Kostović, 2020). Essential for cortical development, these neurons are also responsible for early cortical activity following sensory simulation, and perturbations of these neurons play a role in the development of many neurodevelopmental disorders (Kanold, 2009; Kostović, 2020; Leikos, Tokariev, Koolen, Nevalainen, & Vanhatalo, 2020; Molnár, Luhmann, & Kanold, 2020). Overall, the result of these maturational processes is expressed in EEG by a decrease in discontinuity, an increase in the duration of bursts, a decrease in amplitude, and an increase in the frequency of the EEG signal. At the same time, the content of the bursts evolves with the sequential appearance of EEG patterns during the 3^rd^ trimester such as the TTA-SW, the delta brushes which correspond to oscillators of different frequencies, locations and functionalities (see Wallois et al. (2021) for a review). These dynamics observed during the 3^rd^ trimester constitute the substrate for the complexification of neural interactions which, for example, allow the coding and differentiation of voices, phonemes, and tones (Mahmoudzadeh et al., 2013; Mahmoudzadeh, Wallois, Kongolo, Goudjil, & Dehaene-Lambertz, 2017; Moser et al., 2020; Draganova et al., 2008; Draganova et al., 2007; Sheridan et al., 2008; Hartkopf et al., 2016) by the solicitation of information-coding procedures which, albeit certainly partially immature, are nonetheless completely functional.

This increasing complexity is also expressed by functional connectivity which indicates that the neuronal system appears to evolve from a somewhat random network into a more ordered, “small-world” network (Betzel et al., 2014; Omidvarnia et al., 2014; Tóth et al., 2017). This is in line with the relative immaturity of direct anatomical connections between remote cortical sites at this age (Dubois et al., 2008), with inter- and intra-hemispheric interactions facilitated during the occurrence of bursts which increase during GA.

Overall, the increasing complexity of brain signals, quantified by MSE, might reflect the integration and coordination of neural activity across different spatial or temporal scales. Consequently, the increasing MSE values indicate the development of more organized and complex brain functioning as the premature infants approach full-term gestation.

The MSE-HbR and MSE-TOI parameters, derived from the fNIRS recordings, decrease over GA. MSE-HbO and MSE-HbD also tend to decrease over GA but, in this case, the p-values do not survive correction for multiple comparisons. Since we are not aware of any previous studies on the development of fNIRS signal complexity in infants, we propose that a decrease of MSE over GA may reflect a more efficient and regulated blood supply to the developing brain, indicating improved vascular maturation and oxygenation as gestational age progresses. Alternatively, the increase in the duration of SATs and their decrease in amplitude in the course of gestation might lead to a less marked, and therefore less chaotic NVC. Furthermore, the various NVC strategies reported by Nourhashemi et al. (2020) probably combine efforts towards the end of the last trimester to produce a more robust and less variable vascular response to neural activation. This theory can be linked to the development of cerebral autoregulation during early stages of infancy (Thewissen et al., 2018). In accordance with the rapid development of the human brain in the final trimester of gestation, and with the increasing occurrence of SATs, the decrease of MSE in fNIRS signals might then be the result of a more complex brain activation with less marked NVC. Another explanation could lie in reports of an inverse correlation between oxygen saturation and sample entropy (Bhogal & Mani, 2017). In fact, with the emergence of respiratory movements (Gustafson, May, Yeh, Million, & Allen, 2012) and consequent increase of respiratory capacity in the last trimester of gestation, older preterm infants might have a higher and more stable oxygen saturation which leads to a lower MSE. This interpretation is also in agreement with reports of a positive relationship between cerebral oxygenation and GA at birth in premature infants (Dix, van Bel, & Lemmers, 2017).

Relative to PSD, we observed no changes in the course of GA for the proportion-beta, proportion-delta and for proportion-theta, and a marginally significant increase for proportion-alpha (*Supplementary figure 2*). This is partially in line with previous reports on the maturational changes in the power spectrum in preterm infants (Cohen et al., 2018; Niemarkt et al., 2011; Wel et al., 2017). However, unlike these publications, which described a decrease in the proportion of power in the delta band, we observed no changes during GA. A decrease in TTA-SW and an increase in delta brushes comprised of theta-or gamma-nested activity in a slow wave should also be considered.

Finally, the increase of proportion-SAT during GA was also to be expected, since brain maturation and growth is related to a decrease in quiescent periods, and to an increase in burst duration which ends by the trace alternant in quiet sleep and continuous EEG in active sleep (Benders et al., 2015; Wallois et al., 2021).

### 4.2. Association between MSE and brain activity during GA

The stepwise regression analyses (*Table 2*) revealed several important findings. MSE-EEG is dependent on GA and on proportion-beta. The role of proportion-beta in determining MSE-EEG is also supported in the mediation analysis (*Supplementary figure 3e*), and further explored in *Figure 1f* where we found a positive correlation between proportion-beta and MSE-EEG. Notably, the correlation effect size is high with a Cohen’s *f*^*2*^= 1,24. This relation is probably connected to the beta activity in the delta brushes which increases during the last trimester of gestation (Wallois et al., 2021). The finding that the EEG power in the beta band (13-30 Hz) contributes significantly to MSE suggests that neuronal synchronization within this frequency range plays a crucial role in the maturation of brain networks. Beta band oscillations are associated with various cognitive processes, including attention, sensorimotor integration, and working memory (e.g. Schmidt et al. (2019)). The increasing contribution of beta power to MSE may indicate the emergence of these cognitive functions during the developmental period studied. However, the relationship between power spectrum and complexity metrics is currently a matter of debate; the two parameters may carry shared information for fine-scales, and they may be overlapping representations of neural noise (Kosciessa et al., 2020; Miskovic, MacDonald, Rhodes, & Cote, 2019). In addition, MSE-HbR is dependent on the EEG signal and, specifically, on the SATs. SATs refer to short-lived bursts of activity in neural networks that are thought to play a role in sculpting and refining neuronal connections during development (e.g. Feller (1999); Omidvarnia et al. (2014)). The relationship between SATs and MSE suggests that the complexity of fNIRS signals may be influenced by these spontaneous activity patterns, providing insights into the dynamic processes underlying brain maturation. Finally, the dependency of MSE-TOI on both brain and hemodynamic signals highlights how cerebral oxygenation during GA is subject to complex dynamics.

### 4.3. Brain activity does not mediate the relation between GA and MSE

In our mediation analyses, we did not find any mediation effect of proportion-SAT, proportion-delta, proportion-theta, proportion-alpha or proportion-beta on the relation between GA and MSE calculated on the EEG and fNIRS signals (*Supplementary figures 3 to 7*). With regard to MSE-HbR (*Supplementary figure 5*), the mediation model did not highlight a dependency of this parameter on proportion-SAT such as was reported in the stepwise regression analysis. This might be due to the difference in how the two statistical models were calculated. In fact, mediation stepwise regression models have a different number of possible predictors which impact the correlation values.

### 4.4. Limitations and outlook

One limitation consists in choosing different data lengths for the EEG and fNIRS segments used for MSE calculation (N = 15360 for EEG and N = 2000 for fNIRS). The choice of these data lengths was necessary on account of the different sampling frequencies of the two technologies and to ensure a proper MSE calculation. This entailed calculating MSE over different time scales in the two recording devices to prevent an interpretation of our results on single events. For example, an examination of signal complexity during a SAT in both our EEG and fNIRS recordings is impossible due to the insufficient number of data-points available during a SAT in the fNIRS recording. Moreover, longer recordings could characterize EEG and fNIRS signal complexity during different times of day, potentially highlighting circadian fluctuations. Additionally, we focused our analysis on a single channel in a frontal position in which the EEG and fNIRS sensors were placed on the same spatial location. A multi-channel recording would facilitate a topographic description of MSE in relation to SAT and PSD. Furthermore, a follow-up clinical evaluation of the neonates involved in our analysis would enable us to determine whether our MSE values have a predictive value for the neurological outcome. The original work by Nourhashemi et al. (2020) is currently the only simultaneous EEG-fNIRS dataset available in preterm infants. Here, the behavioral states (BSs), such as sleep or wake, were not registered. BSs have already been shown to have an impact on MSE and on the power of the different frequency bands in the EEG recordings of premature infants (Janjarasjitt, Scher, & Loparo, 2008b; Knyazev, 2012; Wel et al., 2017) and their classification should therefore be taken in account in future work. This is also relevant considering the relationship between BSs, preterm birth, and brain functional connectivity (Uchitel, Vanhatalo, & Austin, 2022). Finally, additional recordings of global physiological noises and a multichannel CW-fNIRS device would allow a better characterization of the fNIRS signal (e.g. Yücel et al., 2021).

## 5. Conclusions

Our results are the first description of developmental trends of signal complexity of multimodal EEG – fNIRS recordings in preterm infants between 27 and 34 weeks of GA. While in the EEG an increase in MSE during GA could reflect the maturational processes in the brain networks during the last trimester of gestation, the decrease in complexity in fNIRS could indicate a smoothing effect on the NVC. This smoothing is not necessarily related to SATs, since discontinuity in the EEG decreases during gestation, resulting in a less chaotic NVC. We also described how spontaneous electrical activity and power in the EEG frequency bands are related to the complexity metrics. We propose that the proportion of EEG power in the beta band (13-30 Hz) might be the main contributor to MSE in the EEG. We also highlight the importance of SATs in determining MSE in the deoxygenated hemoglobin signal derived from fNIRS recording. In summary, these correlation findings in premature infants provide indications of brain maturation, neuronal synchronization, vascular maturation, and the influence of spontaneous activity transients. These findings contribute to our understanding of the complex and nonlinear dynamics of brain development during the early stages of life. Further research and exploration of these associations can help uncover the intricate mechanisms and processes that underlie healthy and pathological brain development in premature infants. For example, multi-channel devices and follow-up clinical evaluation to clarify whether MSE metrics can be used as predictor for neurological outcomes.

## 6. Acknowledgement

This study was conducted with the support of the Procope Mobility program of the French Embassy in Germany, which is designed to enhance scientific cooperation between France and Germany. The work was also partially funded by the ANR-DFG French-German collaboration (PR 496/11-1). We also thank the International Max Planck Research School for the Mechanisms of Mental Function and Dysfunction (IMPRS-MMFD) and the Joachim Herz Foundation.

## 7. Author contributions

L.S. conceptualized and conducted the analysis, drafted and revised the manuscript, and prepared the figures. M.N. and M.M. performed the original recordings. K.S. assisted with the analysis and drafting of the manuscript. M.M., F.W. and H.P. supervised the work. All authors discussed the results and implications, reviewed and edited the manuscript, and approved its final version.

## 9. Data availability statement

The current publication is based on the data from Nourhashemi et al. (2020). In-house codes used for the analysis are available on the OSF page at https://osf.io/v72pz/.

