## Supplementary Material for "Multiscale entropy analysis of combined EEG-fNIRS measurement in preterm neonates"

**Supplementary materials**

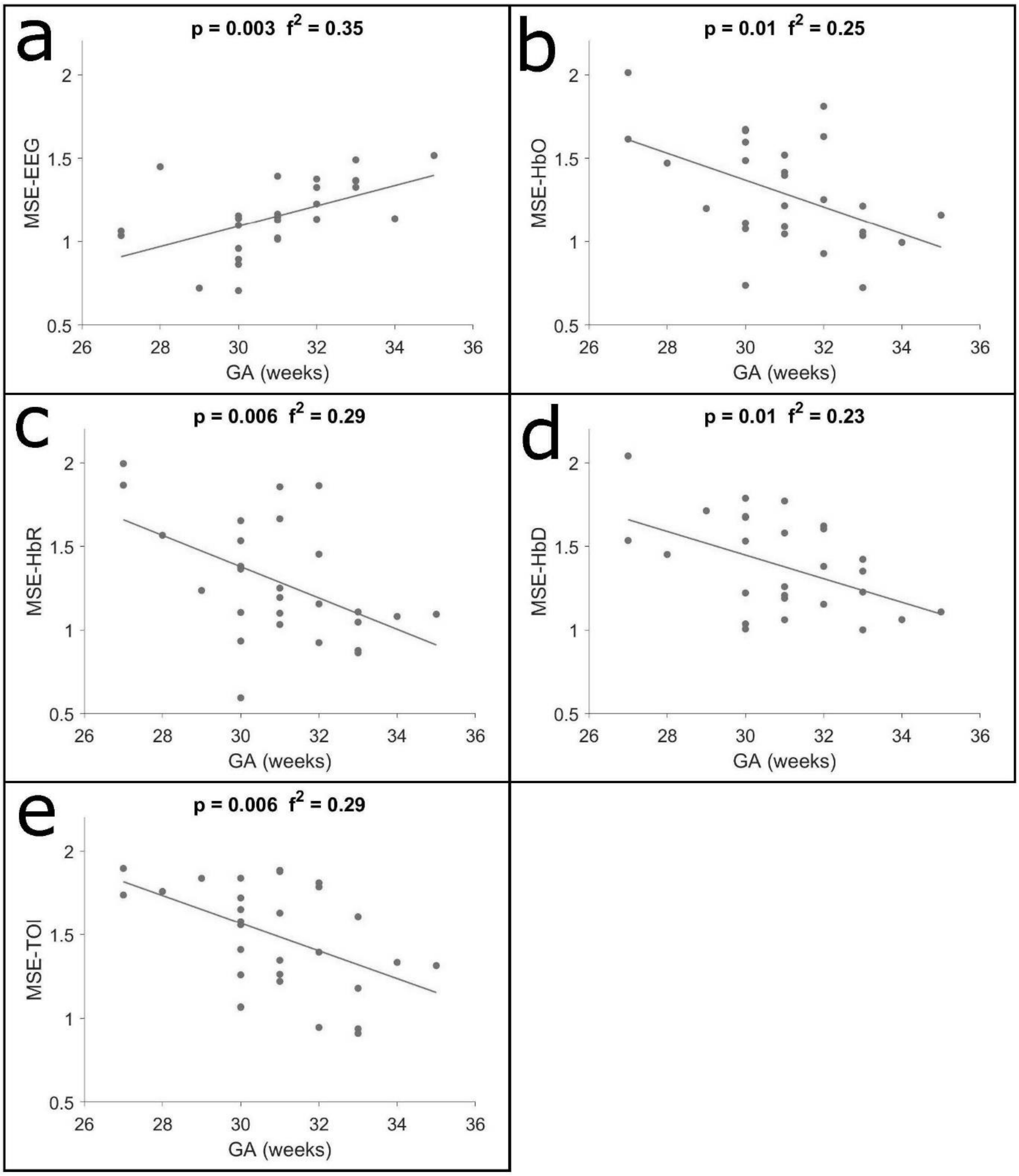


***Supplementary figure 1**. MSE over GA. **a)** increase of MSE-EEG during GA; **b)** probable decrease of MSE-HbO during GA; **c)** decrease of MSE-HbR during GA; **d)** probable decrease of MSE-HbD during GA; **e)** decrease of MSE-TOI during GA.*

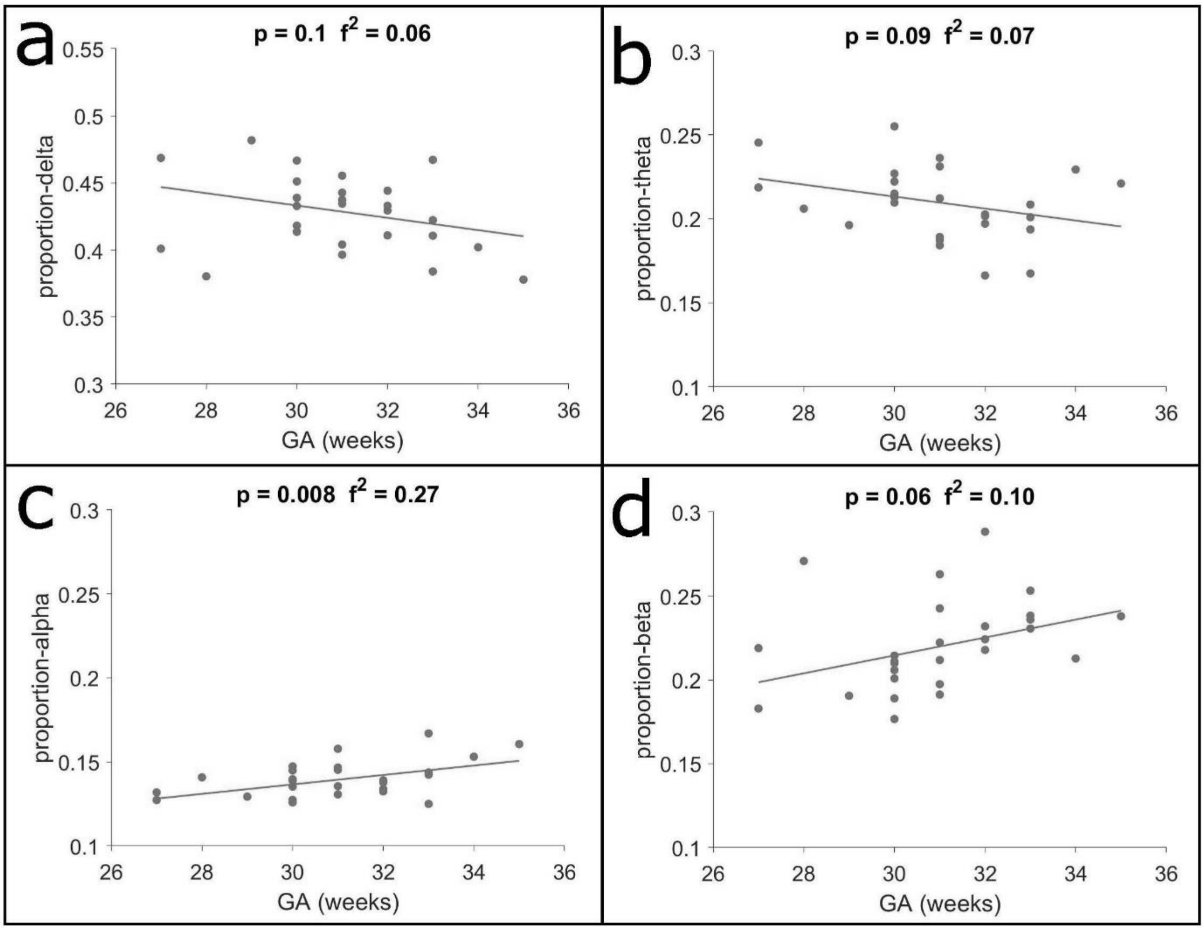


***Supplementary figure 2**. Proportion of PSD during GA. **a)** no changes for proportion-delta during GA; **b)** no changes for proportion-theta during GA; **c)** probable increase of proportion-alpha during GA; **d)** no changes for proportion-beta during GA.*

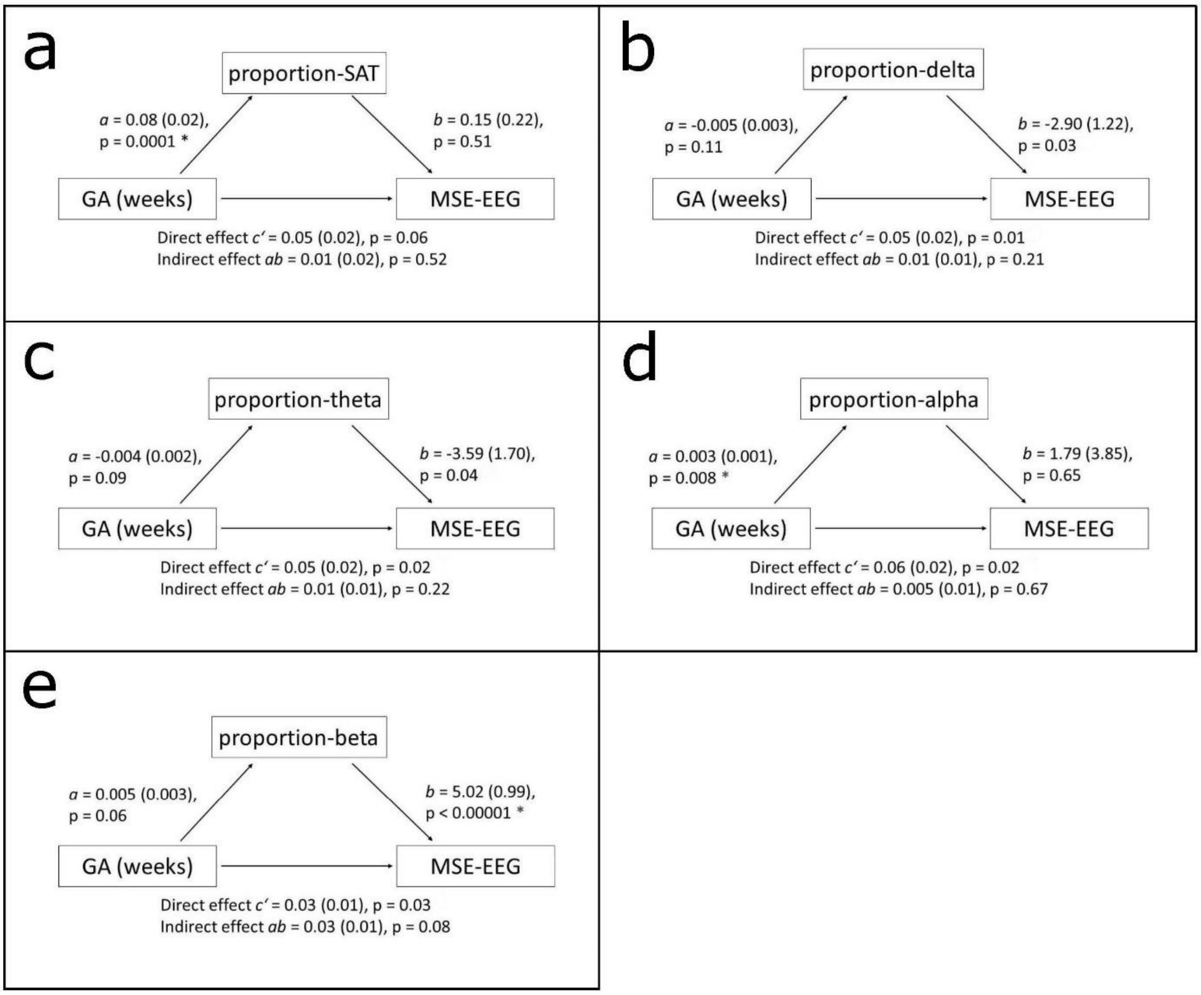


***Supplementary figure 3****. Mediation models with MSE-EEG as outcome variable. * indicates associations which are significant after Holm correction. Path a indicates the relation between GA (weeks) and either proportion-SAT, proportion-delta, proportion-theta, proportion-alpha or proportion-beta; path b indicates the relation between either proportion-SAT, proportion-delta, proportion-theta, proportion-alpha or proportion-beta and MSE-EEG; path c' indicates the direct effect of GA (weeks) on MSE-EEG; path ab indicates the indirect effect of GA (weeks) on MSE-EEG via either proportion-SAT, proportion-delta, proportion-theta, proportion-alpha or proportion-beta.* ***a)*** *GA (weeks) is related to proportion-SAT; proportion-SAT does not mediate the association between GA (weeks) and MSE-EEG.* ***b)*** *Proportion-delta does not mediate the association between GA (weeks) and MSE-EEG.* ***c)*** *proportion-theta does not mediate the association between GA (weeks) and MSE-EEG.* ***d)*** *Proportion-alpha does not mediate the association between GA (weeks) and MSE-EEG.* ***e)*** *Proportion-beta is related to MSE-EEG; proportion-beta does not mediate the association between GA (weeks) and MSE-EEG.*

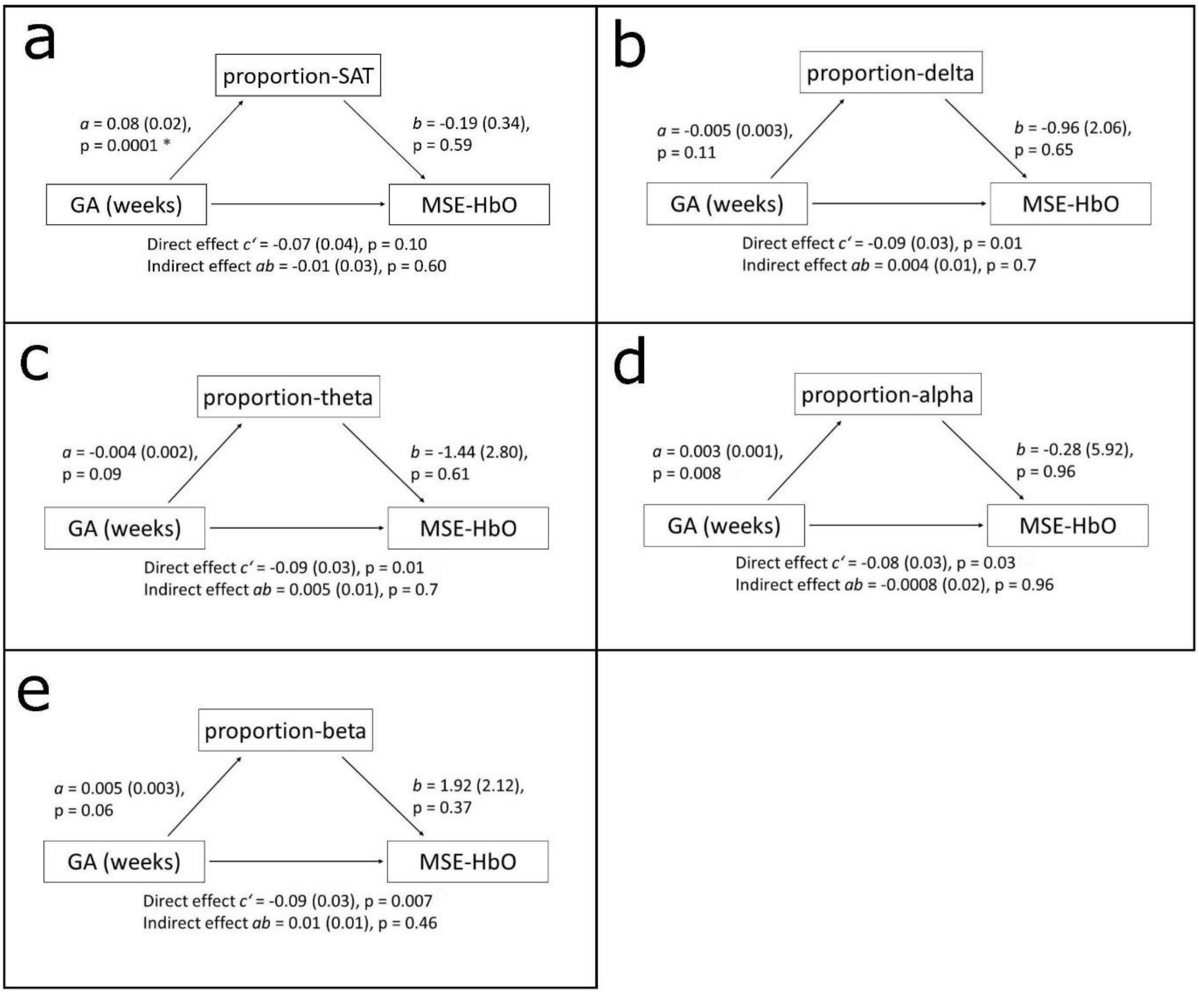


***Supplementary figure 4***. *Mediation models with MSE-HbO as outcome variable. * indicates associations for which p<0.01.* ***a)*** *GA (weeks) is related to proportion-SAT; proportion-SAT does not mediate the association between GA (weeks) and MSE-HbO.* ***b)*** *Proportion-delta does not mediate the association between GA (weeks) and MSE-HbO.* ***c)*** *Proportion-theta does not mediate the association between GA (weeks) and MSE-HbO.* ***d)*** *Proportion-alpha does not mediate the association between GA (weeks) and MSE-HbO.* ***e)*** *Proportion-beta does not mediate the association between GA (weeks) and MSE-HbO.*

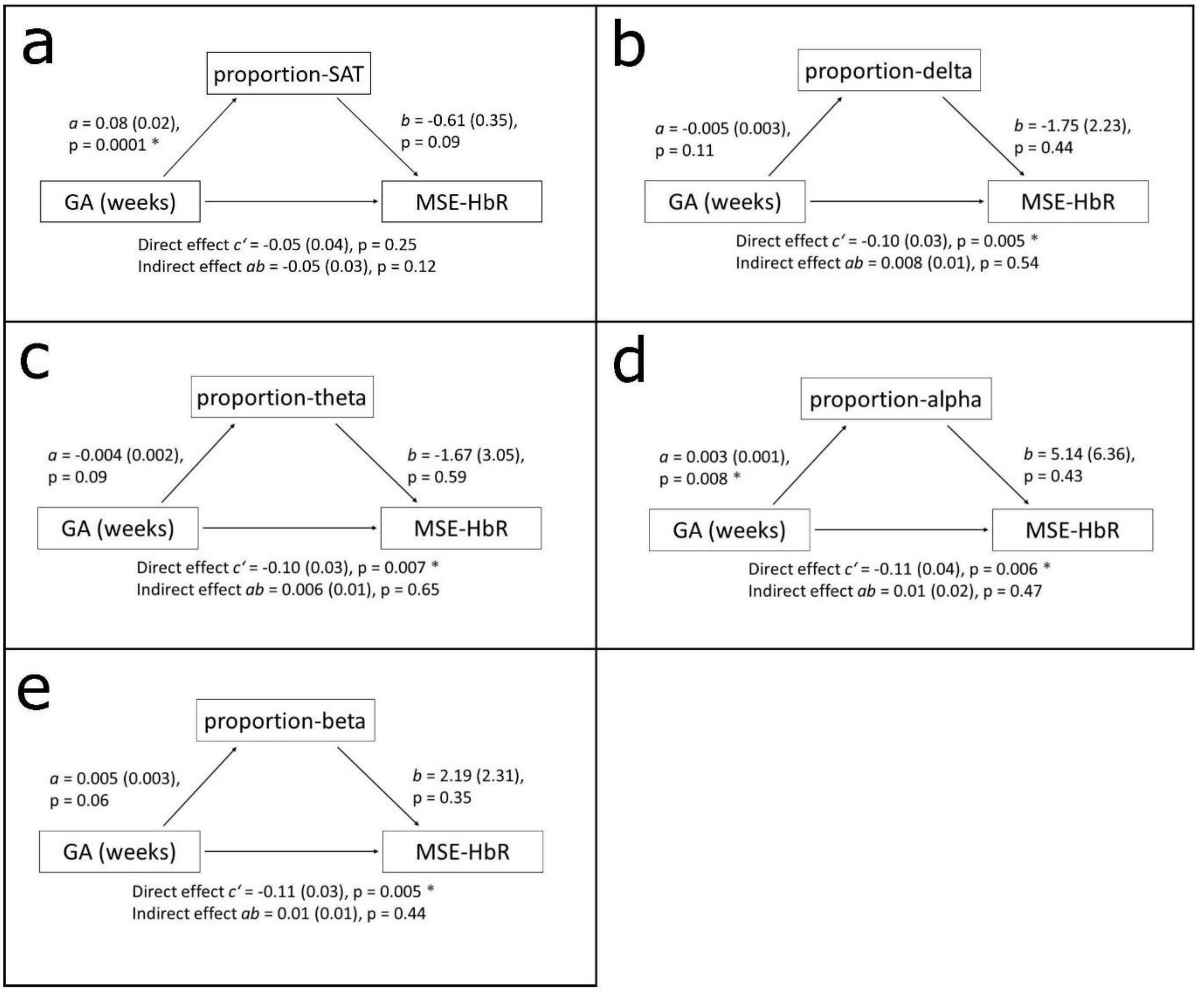


***Supplementary figure 5.*** *Mediation models with MSE-HbR as outcome variable. * indicates associations which are significant after Holm correction.* ***a)*** *GA (weeks) is related to proportion-SAT; proportion-SAT does not mediate the association between GA (weeks) and MSE-HbR.* ***b)*** *GA (weeks) is related to MSE-HbR; proportion-delta does not mediate the association between GA (weeks) and MSE-HbR.* ***c)*** *GA (weeks) is related to MSE-HbR; proportion-theta does not mediate the association between GA (weeks) and MSE-HbR.* ***d)*** *GA (weeks) is related to proportion-alpha; proportion-alpha does not mediate the association between GA (weeks) and MSE-HbR.* ***e)*** *GA (weeks) is related to MSE-HbR; proportion-beta does not mediate the association between GA (weeks) and MSE-HbR.*

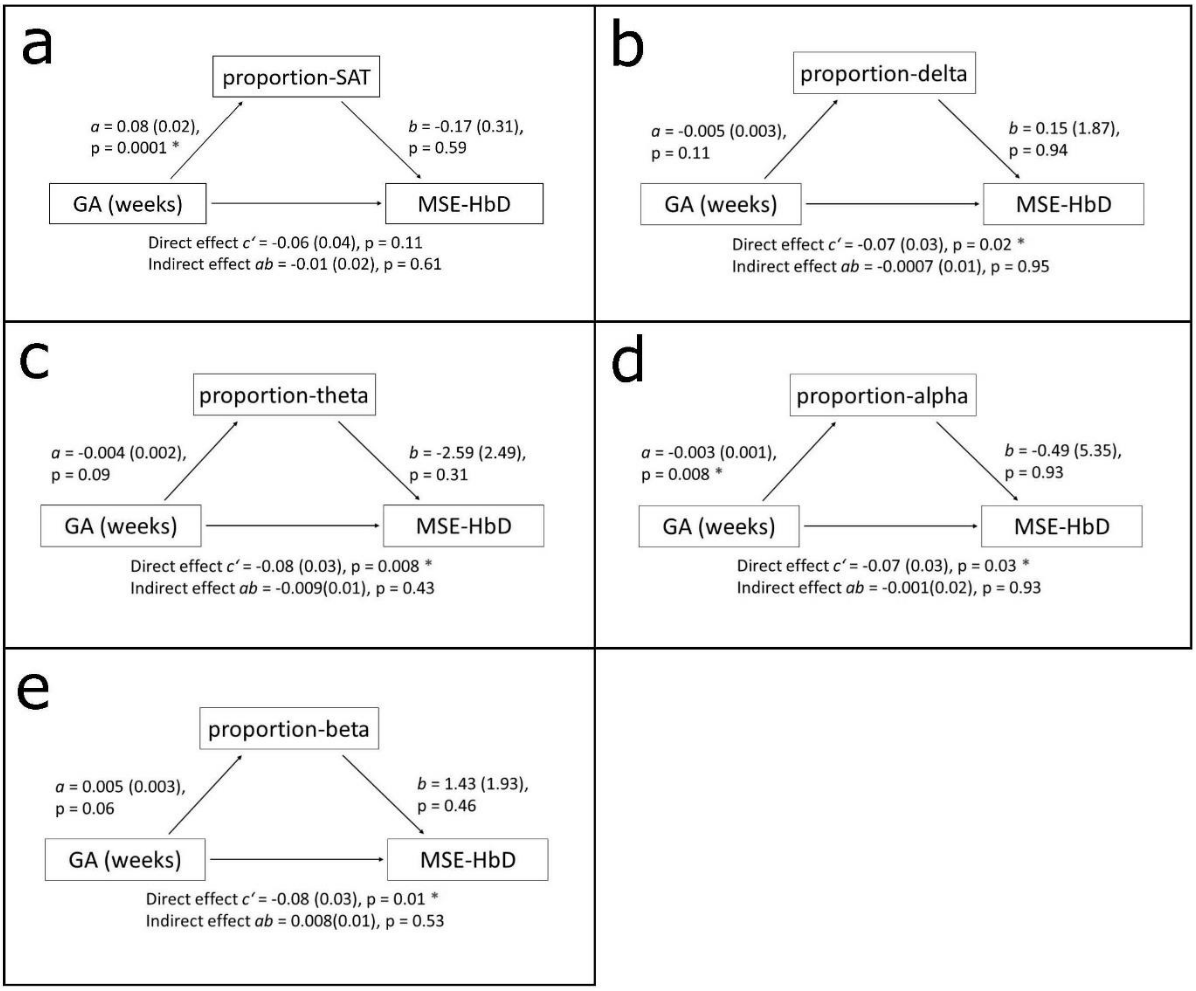


***Supplementary figure 6.*** *Mediation models with MSE-HbD as outcome variable. * indicates associations which are significant after Holm correction.* ***a)*** *GA (weeks) is related to proportion-SAT; proportion-SAT does not mediate the association between GA (weeks) and MSE-HbD.* ***b)*** *proportion-delta does not mediate the association between GA (weeks) and MSE-HbD.* ***c)*** *GA (weeks) is related to MSE-HbD; proportion-theta does not mediate the association between GA (weeks) and MSE-HbD.* ***d)*** *GA (weeks) is related to proportion-alpha; proportion-alpha does not mediate the association between GA (weeks) and MSE-HbD.* ***e)*** *GA (weeks) is related to MSE-HbD; proportion-beta does not mediate the association between GA (weeks) and MSE-HbD.*

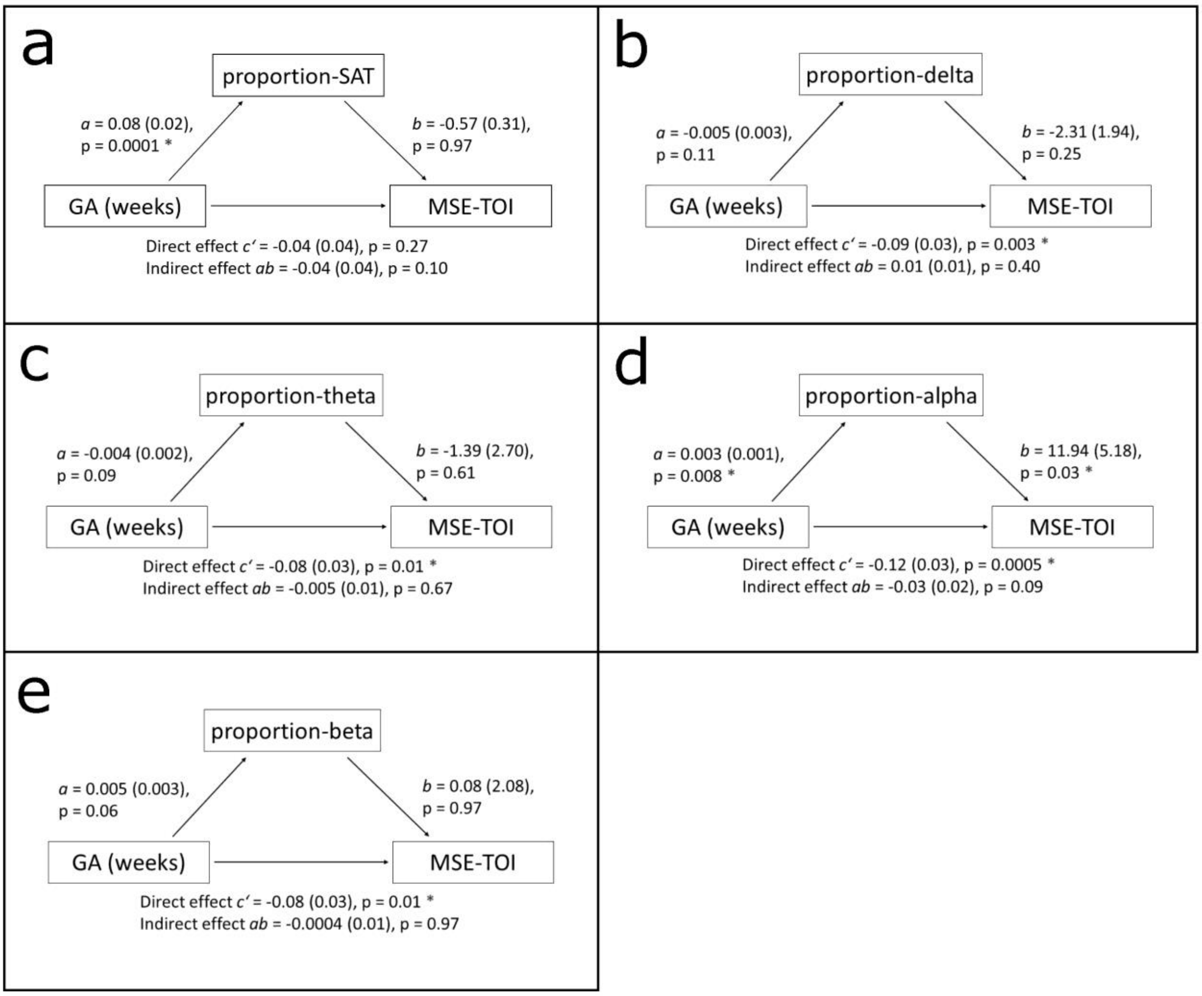


***Supplementary figure 7.*** *Mediation models with MSE-TOI as outcome variable. * indicates associations which are significant after Holm correction.* ***a)*** *GA (weeks) is related to proportion-SAT; proportion-SAT does not mediate the association between GA (weeks) and MSE-TOI.* ***b)*** *GA (weeks) is related to MSE-TOI; proportion-delta does not mediate the association between GA (weeks) and MSE-TOI.* ***c)*** *GA (weeks) is related to MSE-TOI; proportion-theta does not mediate the association between GA (weeks) and MSE-TOI.* ***d)*** *GA (weeks) is related to proportion-alpha and to MSE-TOI; proportion-alpha does not mediate the association between GA (weeks) and MSE-TOI.* ***e)*** *GA (weeks) is related to MSE-TOI; proportion-beta does not mediate the association between GA (weeks) and MSE-TOI.*
